# Elevated developmental temperatures below the lethal limit reduce *Aedes aegypti* fertility

**DOI:** 10.1101/2024.10.10.617713

**Authors:** Miriama Pekľanská, Belinda van Heerwaarden, Ary A. Hoffmann, Marcela Nouzová, Radek Šíma, Perran A. Ross

## Abstract

*Aedes aegypti* mosquitoes are the principal vectors of dengue and continue to pose a threat to human health, with ongoing urbanization, climate change, and trade all impacting the distribution and abundance of this species. Hot periods are becoming increasingly common and their impacts on insect mortality have been well established, but they may have even greater impacts on insect fertility. In this study, we investigated the impacts of high temperatures on *Ae. aegypti* fertility both within and across generations. Mosquitoes developing under elevated temperatures exhibited higher critical thermal maxima (CTmax) reflecting developmental acclimation, but their fertility declined with increasing developmental temperature. In females, elevated developmental temperatures decreased fecundity while in males it tended to decrease egg hatch proportions and the proportion of individuals producing viable offspring. Rearing both sexes at 35°C increased fecundity in the subsequent generation but effects of elevated temperatures persisted across gonotrophic cycles within the same generation. Moreover, exposure of adults to 35°C further decreased fertility beyond the effects of developmental temperature alone. These findings highlight sub-lethal impacts of elevated temperatures on *Ae. aegypti* fertility and plastic responses to thermal stress within and across generations. This has significant implications for mosquito populations thriving in increasingly warmer environments.

## Introduction

Mosquitoes are significant vectors of disease-causing agents such as dengue, Zika, chikungunya, and malaria, presenting a global hazard to public health (Bhatt et al., 2013). As ectotherms, mosquitoes are highly sensitive to temperature fluctuations, making environmental changes particularly impactful (Paaijmans et al., 2013). Temperature fluctuations during their development can be significant, often influenced by the type of water reservoir and its exposure to sunlight (Richardson et al., 2013).

Temperature not only influences mosquito development, survival and reproduction (Agyekum et al., 2021; Eisen et al., 2014), but also their vector competence (Bellone & Failloux, 2020; Carrington, Seifert, Armijos, et al., 2013). Therefore, understanding the upper thermal limits of various traits is essential for predicting how mosquito populations and pathogens they spread will respond to climate change (Couper et al., 2021; Lahondère & Bonizzoni, 2022).

Mosquito development and mortality follow characteristic thermal performance curves, where temperatures above and below the optimal range result in decreased population growth (Eisen et al., 2014), with reduced egg hatching frequencies (Farnesi et al., 2009; Mohammed & Chadee, 2011; Sasmita et al., 2019; Sukiato et al., 2019), reduced suvival of immature stages (Richardson et al., 2011; Tun-Lin et al., 2000) and decreased adult longevity (Marinho et al., 2016). These thermal optima differ between traits and life stages. In the principal dengue vector *Aedes aegypti*, optimal temperatures for development are warmer than those for survival (Carrington, Armijos, et al., 2013; Dennington et al., 2024; Richardson et al., 2011). Temperature effects can be regulated by behavioral responses (Ziegler et al., 2022, 2023), adaptation through heritable genetic changes (Dennington et al., 2024; Ware-Gilmore et al., 2023) or phenotypic plasticity, where an organism produces alternative phenotypes in different environmental conditions (Chevin et al., 2010; Ghalambor et al., 2007; Sgrò et al., 2016). This is evident in natural mosquito populations where thermal tolerance can shift seasonally (Oliveira et al., 2021) and is correlated with local environmental temperatures (Lyberger et al., 2024).

Plastic responses to temperature can be induced and persist both within and across life stages and generations (Steigenga & Fischer, 2007), affecting a broad range of traits. In mosquitoes, warmer temperatures increase the rate of egg production (Delatte et al., 2009; Rúa et al., 2005) and the develeopmont of embryos (Farnesi et al., 2009; Marinho et al., 2016) and larvae (Couret et al., 2014; Marinho et al., 2016; Sasmita et al., 2019), but reduce adult body size (Farjana et al., 2012; Mohammed & Chadee, 2011) and fecundity (Marinho et al., 2016). These plastic responses can provide benefits in some environments, where acclimation of mosquitoes to hot or cold conditions improves performance in similar conditions as measured by critical thermal maximum (CTmax) or survival assays (Gray, 2013; Ioannou et al., 2024; Jass et al., 2019; Lyons et al., 2012; Mellanby, 1960; Sasmita et al., 2019; Sivan et al., 2020). However, plasticity can also have costs that decrease fitness, particularly when responses are irreversible (Hoffmann & Bridle, 2022). Diapause is one such irreversible change that can be costly if triggered at an inappropriate time (Lacour et al., 2015; Lee & Duvall, 2022). For container-breeding mosquitoes developing in fluctuating thermal environments, inappropriate plastic responses may occur, where the potential benefits are diminished if environmental conditions shift, and fitness may decline due to accumulated heat stress.

Fertility is an important component of individual fitness, and serves as a key predictor of population growth and survival. Recent studies suggest that a species’ ’thermal fertility limit’ (TFL) is the best predictor of a species distribution and their response to climate change (Parratt et al., 2021; Van Heerwaarden & Sgrò, 2021; Walsh et al., 2019). While previous studies have investigated thermal performance curves of *Ae. aegypti* fertility (Carrington, Armijos, et al., 2013; Dennington et al., 2024; Marinho et al., 2016; Yang et al., 2009), these typically only consider egg counts and not egg hatch proportions, potentially underestimating the impacts of temperature on overall fertility. For example, Costa et al. (2010) found that both fecundity and egg hatch decline in female *Ae. aegypti* exposed to elevated temperatures after blood feeding. They also did not consider the cumulative effects of heat stress across life stages, or consider each life stage individually. Temperatures during both development and adulthood can influence fertility, with cumulative costs to fecundity observed in *Ae. albopictus* when both larvae and adults are exposed to elevated temperatures (Ezeakacha & Yee, 2019). Temperature exposure may also differ substantially between pre-adult and adult stages, as they occupy different habitats and vary in their ability to thermoregulate. Costs of elevated temperatures may differ between sexes (Janowitz & Fischer, 2011; Sales et al., 2018; Van Heerwaarden & Sgrò, 2021; Walsh et al., 2021; Zwoinska et al., 2020), but no studies in mosquitoes have considered impacts on the fertility of both males and females. Furthermore, the persistence of these effects across gonotrophic cycles or generations remains underexplored (Weaving et al., 2024) and has not been tested in mosquitoes (Couper et al., 2021).

In this study, we assess the effects of elevated developmental and adult temperatures on *Ae. aegypti* fertility and reproductive success, examining both fecundity and egg hatch proportions as well as the proportion of individuals with viable offspring. We demonstrate that elevated developmental temperatures affect fertility below lethal limits, impacting both females and males in different ways. These effects persist across gonotrophic cycles and are more severe when both immature and adult stages experience elevated temperatures. Additionally, we highlight potential adaptive responses of *Ae. aegypti* to elevated developmental temperatures, including an increase in CTmax and improved fecundity in the subsequent generation. Our findings underscore the importance of considering sub-lethal effects of high temperature on fertility and plasticity when predicting mosquito responses to climate change.

## Material and methods

### Experimental population and maintenance

The *Ae. aegypti* population used here was originally collected from Cairns, Queensland, Australia in 2018 and was free of *Wolbachia* infection. Adults were maintained at a census size of 450-500 individuals at 26°C under a 12:12 light:dark cycle and provided with 10% sucrose solution through soaked cotton balls. Larvae were reared at a controlled density (500 larvae in 4L of reverse osmosis water) and fed fish food (Hikari tropical sinking wafers, Kyorin food, Himeji, Japan). To obtain eggs, females were blood fed on the forearm of a single adult human volunteer which was approved by the University of Melbourne Human Ethics committee (project ID 28583).

### Upper developmental lethal thermal limits

To determine upper developmental lethal thermal limits in this population, *Ae. aegypti* were hatched and reared at temperatures of 26, 28, 30, 32, 34, 35, 36 and 37°C in climate controlled incubators (PHCbi, MIR-254). Eggs (< 2 weeks old) collected on sandpaper (Norton Master Painters P80; Saint-Gobain Abrasives Pty. Ltd., Thomastown, Victoria, Australia) and maintained at 26°C were separated into batches of 50-100 eggs and hatched in 750 mL plastic trays filled with 500 mL of reverse osmosis water that was pre-heated to each temperature. A few grains of yeast were added to each tray to stimulate hatching. One day after hatching, five to six replicate batches of eggs per temperature were measured for their hatch proportion by counting the number of unhatched (intact) and hatched eggs with a clearly detached cap. Emerging larvae were transferred to trays with 500 mL of water with five replicates of 50 larvae per temperature. Larvae were provided with fish food *ad libitum* and maintained at the same temperature throughout the aquatic phase until adult emergence. Larva to pupa viability was calculated by counting the number of individuals reaching pupation and dividing by the number of larvae added to the tray. Pupa to adult viability was calculated by dividing the number of adults by the number of pupae. Adults that did not fully eclose from the pupal case were counted as dead pupae. We also estimated egg to adult viability by multiplying mean egg hatch proportions by the proportion of larvae reaching adulthood at each temperature.

### Effects of elevated developmental temperatures on CTmax

We performed critical thermal maxima (CTmax) assays to measure the effects of elevated developmental temperatures on adult heat tolerance. Mosquitoes were reared as described above from 1st instar larvae to the adult stage at the following constant temperatures: 26, 32, 33, 34 and 35°C. These thermal regimes were chosen as 26°C is within the optimal range of immature survival for Australian populations of *Ae. aegypti* (Richardson et al., 2011; Tun-Lin et al., 2000) and 35 °C was the highest temperature where egg to adult viability was greater than 0.5 in the previous experiment. Adults (2-4 d old, mated) were aspirated into numbered glass vials (50mm height x 18 mm diameter, 10mL volume), sealed with a plastic cap and fastened to a plastic rack with clips in a randomized order. The rack was submerged in a water tank (Ratek Instruments, Boronia, Victoria, Australia) which was programmed to increase in temperature from 25°C at a rate of 0.1°C/min. The water temperature in the middle of the rack was measured with a real-time temperature probe. Mosquitoes in vials were observed and the temperature where an individual ceased movement was recorded to the nearest 0.1°C. We tested 48 replicate individuals per sex and developmental temperature.

### Effects of elevated developmental temperatures on fertility

We measured the impact of a range of sub-lethal temperatures (where egg to adult viability was greater than 0.5) on mosquito fecundity, egg hatch and total offpring counts to determine whether fertility is affected below the lethal limit. Mosquitoes were reared as described above at the following constant temperatures: 26, 32, 33, 34 and 35°C. Pupae were sorted by sex and maintained at the same water temperature until reaching adulthood. All adults were transferred to 26 °C within 2 d post-emergence and maintained for a further 2 d to ensure that they had reached sexual maturity. We then performed crosses at 26°C between males and females reared at the same temperature (26, 32, 33, 34 and 35°C). Mosquitoes were crossed in groups rather than in mating pairs due to space constraints, with cages > 1L in volume needed to ensure high insemination frequencies.

Once crosses were set up, adults were provided with water and a 10% sucrose solution until 24 hr before blood feeding. Females (4-6 d old) were blood fed and 30 females per cross were transfered individually to 70 mL specimen cups containing 20 mL of larval rearing water and lined with a strip of sandpaper to encourage oviposition. Sandpaper strips were collected 4 d after blood feeding, partially dried, then maintained at a high humidity. After 3 d, eggs were hatched at 26 °C and left for 24 hr. Fecundity was determined by counting the total number of eggs per female, with egg hatch proportions calculated by dividing the number of hatched eggs by the total number of eggs. Total offspring per female was determined by counting the total number of hatched eggs. Females that died before laying eggs were excluded from the analysis.

### Sex-specific effects of elevated developmental temperatures on reproductive success and fertility

To investigate sex-specific effects of elevated developmental temperatures, we set up crosses between mosquitoes reared at 26 and 35°C. Mosquitoes were reared as described above at constant temperatures of 26 and 35°C then transferred to 26°C prior to mating. Males (2-4 d old) reared at 26 or 35°C were crossed to females (2-4 d old) reared at 26 or 35°C for a total of four crosses. Each cross involved three replicate cages of 50 mosquitoes per sex. Females (4-6 d old) were blood fed and 30 females per replicate cage were isolated for oviposition. Eggs were collected 4 d after blood feeding then hatched 3 d after collection at 26 °C to measure fecundity, hatch proportions and total offspring as described above. Females that died before laying eggs were excluded from the analysis.

### Combined effects of elevated developmental and adult temperatures on reproductive success and fertility

We measured the combined effects of elevated developmental and adult temperatures on fertility by rearing mosquitoes at 26 and 35°C then maintaining adults at both temperatures. Mosquitoes were reared as described above at 26 and 35°C then 1-2 d old adults were transferred to cages at 26 and 35°C for mating. This gave a total of four treatments, where mosquitoes developed at 26 or 35°C and were then maintained at 26 or 35°C during adulthood, which included mating, blood feeding and egg laying. Both the males and females within a treatment were maintained at the same temperature.

Mosquitoes (4-6 d old) were blood fed and isolated for oviposition, with eggs immediately transferred to 26°C following collection. Females were then measured for their fecundity, hatch proportions and total offspring as described above. We set up 30 replicates for the treatments in which the adults were kept at 26 °C and 60 replicates for the treatments in which the adults were kept at 35 °C, as we expected higher mortality based on pilot experiments.

### Effects of elevated developmental temperatures on reproductive success and fertility across generations and gonotrophic cycles

To measure the effects of elevated developmental temperatures on fertility across generations and gonotrophic cycles, eggs were collected from populations where both sexes developed at 26 or 35°C as described above. Larvae hatched at 26°C were immediately transferred to trays at 26 or 35°C and reared to adulthood. This gave a total of four treatments, where mosquitoes developed at 26 or 35°C in one or both generations. In treatments where mosquitoes developed at 35°C, only larvae and pupae were exposed, with adults and eggs maintained at 26°C. Crosses between males and females within the same treatment were performed as described above with two replicate cages of 50 mosquitoes per sex. Mosquitoes (4-6 d old) were blood fed and 30 females per replicate cage were isolated for oviposition. Eggs were collected 4 d after blood feeding and the sandpaper was replaced. Females were then blood fed individually to initiate a second gonotrophic cycle and these eggs were collected 4 d after the second blood meal. Eggs from both gonotrophic cycles were hatched 3 d after collection at 26 °C to measure fecundity, hatch proportions and total offspring as described above. Females that died before laying eggs were excluded from the analysis.

### Statistical analysis

Statistical analysis for all experiments was conducted using R in RStudio (version 1.3.959). Upper lethal thermal limits (LT_50_) for egg hatchability, larva to pupa viability and pupa to adult viability were calculated using a dose–response model (drm) that was created using the *drc* package (version 3.0-1; (Ritz et al., 2015)).

To examine the impact of developmental temperature and sex on CTmax, we used a two-way ANOVA (*car* package 3.0-12; (Fox & Weisberg, 2018)) on a linear model, with family set as Gaussian, as this distribution fit the data best after exploration using the *DHARMa* package (version 0.4.5; (Hartig, 2018)). We then explored plasticity in CTmax by calculating the acclimation response ratio (ARR) between developmental temperatures of 26 and 35°C and between 32 and 35°C, which estimates the degree change in CTmax as a proportion of difference in developmental temperature (Gunderson & Stillman, 2015; Semsar-kazerouni & Verberk, 2018).

In the fertility experiments, we analysed fecundity (number of eggs laid), egg hatch proportions of those eggs, and total offspring (number or eggs that hatched) as an overall estimate of fertility. For these traits which did not follow a dose response curve across the range of developmental temperatures from 32-35°C, we looked at the effect of temperature using a one-way ANOVA (*car* package 3.0-12) on a linear model, with family set as Gaussian, as this distribution fit the data best after exploration using the *DHARMa* package. Females that died before reproducing were excluded from the analysis as we could not accurately determine whether females died due to impacts of high temperatures or handling, though this led to variable levels of replication between treatments. Females that did not lay eggs were excluded from analyses of egg hatch proportions but included in fecundity and total offspring analyses. Some females laid eggs which could not easily be collected (e.g. some eggs became submerged); these individuals were included in analyses of fecundity but not the other traits.

To assess the effect of male and female developmental temperature (fixed effects: 26 or 35°C) on fecundity, hatch proportion and total offspring, we first explored the effect of developmental temperature and sex on reproductive success (i.e. whether females did or did not lay eggs, had any eggs hatching, or produced any offspring), based on a binomial model with a logit link function. We then tested for effects of developmental temperature and sex on fertility (counts) excluding zero values (where females laid no eggs, had zero eggs hatching or produced no offspring) using a two-way ANOVA (*car* package 3.0-12) on a linear model, with the distribution set as gaussian, as this distribution fit the data best after exploration using the *DHARMa* package (version 0.4.5). To test the effects of elevated developmental and adult temperatures, we used the same sets of analysis but with life stage and temperature as fixed effects. The same analyses were also performed to test for cross generational effects of developmental temperature, with generation (parental and offspring) and developmental temperature (26 or 35°C) as fixed effects, for each gonotrophic cycle.

## Results

### Developmental lethal thermal limits differ between life stages

We reared *Ae. aegypti* at a range of temperatures to determine the developmental lethal thermal limits at each life stage (Figure 1). Eggs hatched well in water temperatures up to 36°C but experienced a sharp decline in hatch proportion at 37°C, with a LT_50_ of 36.51°C (lower, upper 95% confidence interval: 36.38, 36.65). Larvae experienced a similar decline in survivorship to the pupal stage with a LT_50_ of 36.34°C (36.26, 36.43). Pupae were even more sensitive to elevated temperatures, with a LT_50_ of 35.70°C (35.58, 35.82) and zero pupae reaching adulthood at 37°C (Figure 1), which may reflect heat damage accumulated during both the larval and pupal stage. When considering viability across all immature stages (from egg hatching to adult emergence), fewer than 50% of individuals survived to the adult stage at 36°C (Figure 1). To minimize potential effects of selection, we conducted subsequent experiments at constant temperatures of 35°C and below.

**Figure 1.**
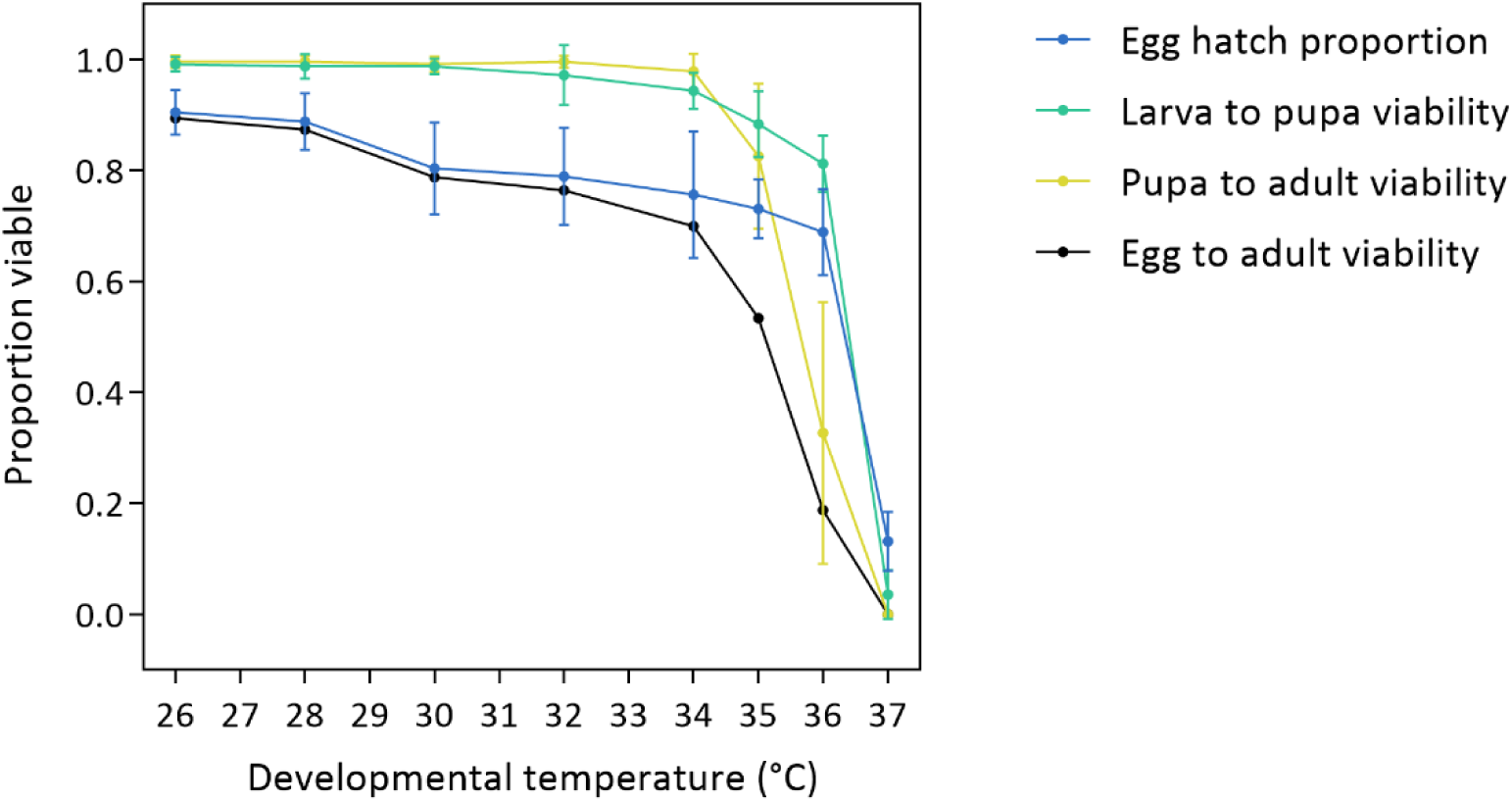
Developmental lethal thermal limits of *Aedes aegypti* at constant temperatures. We measured the proportion of eggs (blue), larvae (green) and pupae (yellow) reaching the next life stage at a range of constant temperatures. Dots and error bars represent means and 95% confidence intervals based on 5-6 replicate batches of eggs for hatch proportions and 5 replicate trays of 50 larvae for larva to pupa and pupa to adult viability. Egg to adult viability (black) was estimated by multiplying mean egg hatch proportions by the proportion of larvae reaching adulthood at each temperature.

### Developmental temperatures below the lethal limit increase CTmax

We assessed the upper thermal limits of adult *Ae. aegypti* reared at elevated developmental temperatures by measuring their CTmax. We found a significant effect of developmental temperature (two-way ANOVA: F_1,524_ = 61.973, P < 0.001), with CTmax increasing as developmental temperature rose for both sexes (Figure 2). There was no significant effect of sex (F_1,524_ = 2.587, P = 0.108) nor was there a significant interaction between developmental temperature and sex (F_1,524_ = 1.867, P = 0.172). When comparing the two most extreme developmental temperatures (26 and 35°C), we calculated an ARR of +0.037°C per °C of developmental temperature for females and +0.026°C for males. This increased to +0.062°C for females and 0.047°C for males when comparing the ARR between developmental temperatures of 32 and 35°C. These results demonstrate developmental plasticity in CTmax for both sexes, with elevated developmental temperatures leading to higher CTmax values.

**Figure 2.**
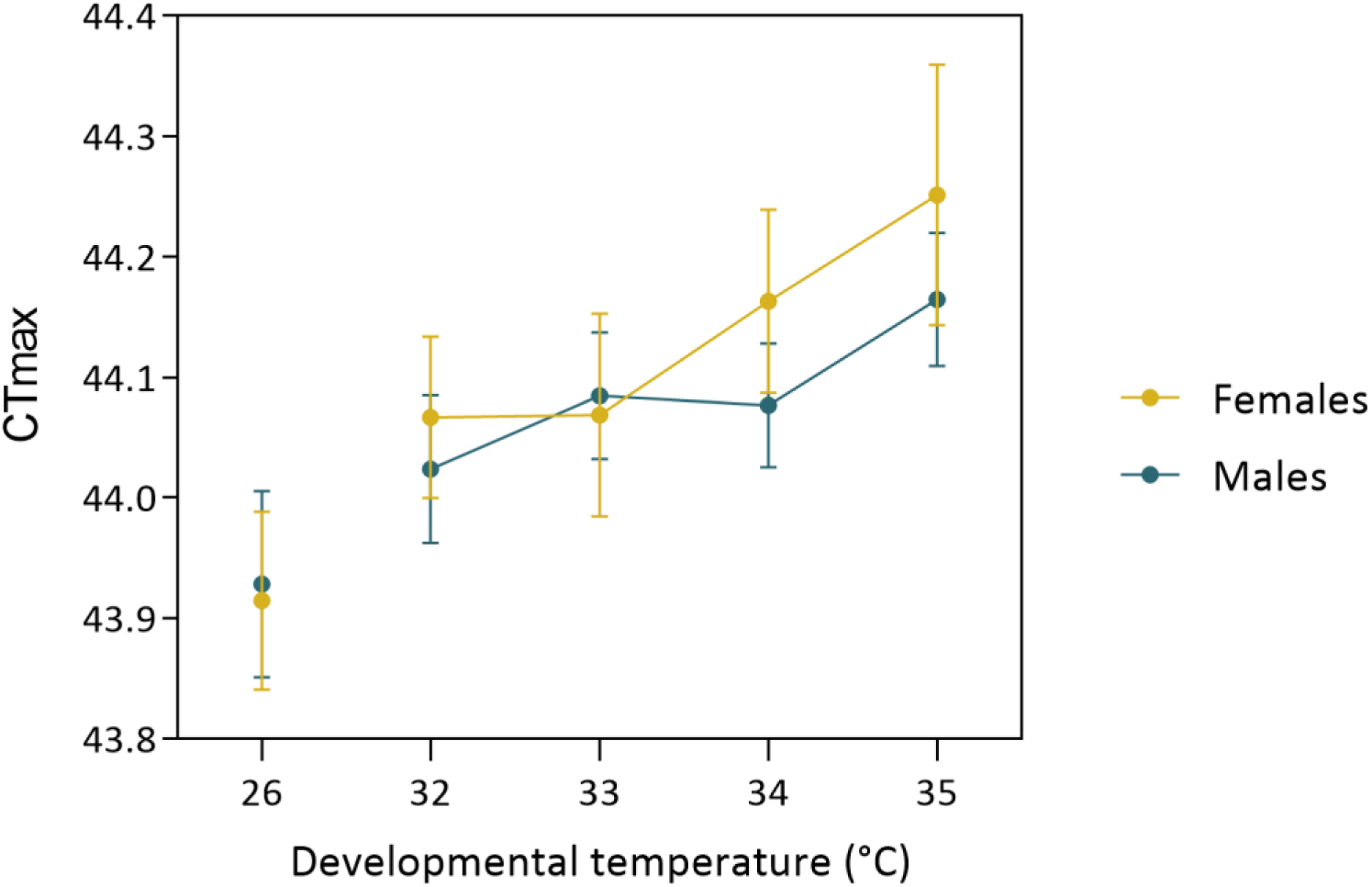
CTmax of *Aedes aegypti* females (yellow) and males (blue) reared at sub-lethal elevated temperatures. Dots and error bars represent means and 95% confidence intervals based on 48 replicates (females) or 60 replicates (males) per developmental temperature.

### Fertility declines below the lethal developmental thermal limit

We reared *Ae. aegypti* at control (26°C) or elevated (32°C, 33°C, 34°C and 35°C) temperatures from egg to adult to assess the impact of sub-lethal developmental temperatures on their fertility. We observed significant effects of temperature on fecundity (one-way ANOVA: F_1,107_ = 6.746, P = 0.011), egg hatch proportion (F_1,98_ = 17.881, P < 0.001) and total offspring (F_1,103_ = 16.596, P < 0.001), with higher developmental temperatures leading to a decrease in fertility (Figure 3). Although fertility declined, mosquitoes remained fertile at 35°C and there was no further significant decrease beyond 33-34°C across all traits (Figure 5).

**Figure 3.**
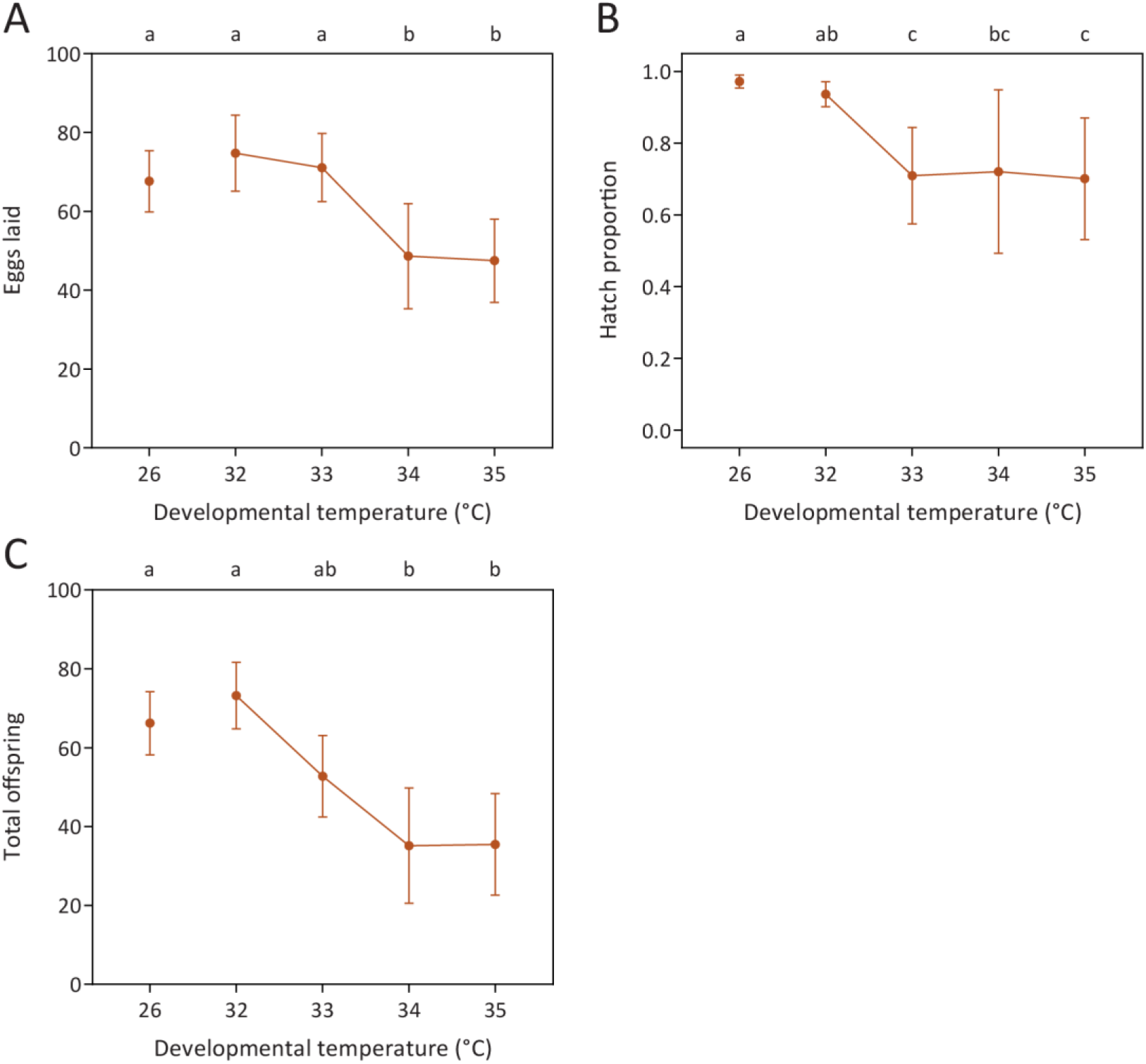
Fertility of *Aedes aegypti* reared at sub-lethal elevated temperatures. We measured (A) fecundity, (B) egg hatch proportions and (C) total offspring when both sexes developed at elevated temperatures. Dots and error bars represent medians and 95% confidence intervals based on up to 30 replicates per cross and temperature (females that died prior to egg laying were excluded from the analysis). Different letters above plots indicate significant differences (P < 0.05) between treatments according to Tukey’s post-hoc tests.

### Effects of elevated developmental temperatures on reproductive success and fertility are sex-specific

To test for sex-specific effects of elevated developmental temperatures on reproductive success and overall fertility, we peformed crosses where males, females or both sexes developed at either 26 or 35°C. Rearing males at 35°C decreased their reproductive success (Figure 4), with an analysis of the proportional data indicating substantial effects of male rearing temperature on the proportion of females laying eggs (two-way ANOVA: χ^2^ = 35.544, df = 1, P < 0.001), the proportion of females with some viable eggs (χ^2^ = 26.983, df = 1, P < 0.001), and the proportion of females with some viable offspring (χ^2^ = 37.824, df = 1, P < 0.001). While there were no significant effects of female rearing temperature on these traits (all P > 0.114), we found sigificant interactions between male and female rearing temperature on the proportion of females laying eggs (χ^2^ = 8.771, df = 1, P = 0.003) and the proportion of females with some viable offspring (χ^2^ = 9.081, df = 1, P = 0.003) which was driven by a relatively low proportion of females in the 26 ♀ x 35 ♂ cross that laid eggs.

**Figure 4.**
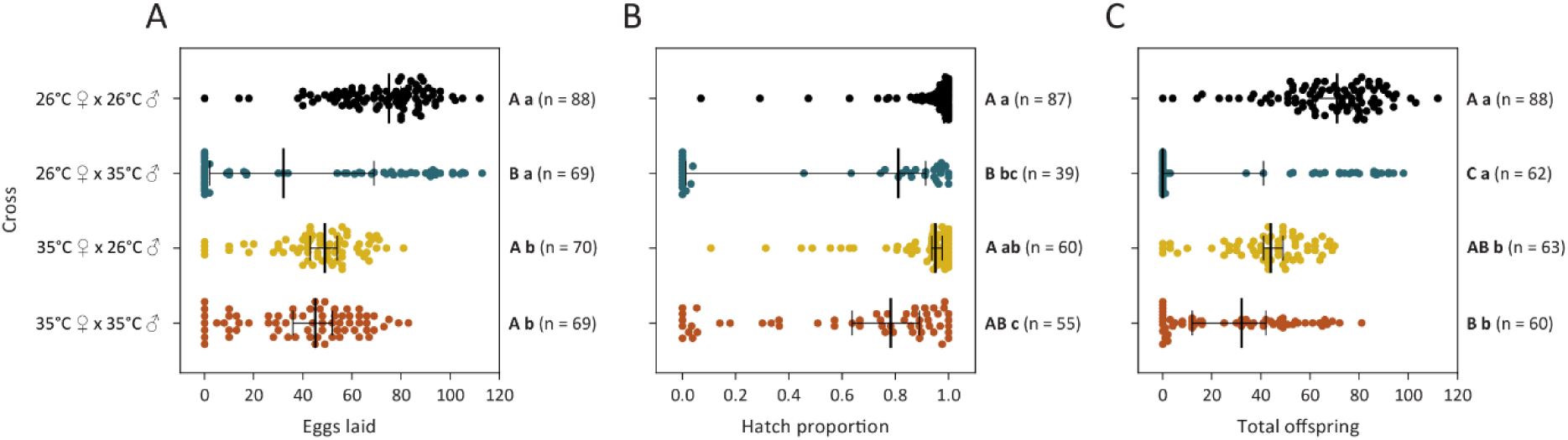
Fertility and reproductive success of *Aedes aegypti* females in crosses between males and females reared at 26 or 35°C. We measured (A) fecundity, (B) egg hatch proportions and (C) total offspring. Dots and error bars represent medians and 95% confidence intervals. Different letters next to plots indicate significant differences (P < 0.05) between treatments according to Tukey’s post-hoc tests, with capital letters for the binomial analyses of reproductive success and lowercase letters for the analyses of counts excluding zero values. The n values represent the number of replicates measured per treatment and trait.

When considering counts with zero values excluded, females reared at 35°C had significantly reduced fecundity (Two-way ANOVA: F_1,254_ = 65.883, P < 0.001) and total offspring (F_1,221_ = 78.891, P < 0.001) but not egg hatch proportions (F_1,221_ = 3.895, P = 0.050, Figure 4). Male developmental temperature also signifcantly affected fecundity (F_1,254_ = 4.023, P = 0.046) and total offspring (F_1,221_ = 4.025, P = 0.046) but had the most substantial effect on egg hatch proportion (F_1,221_ = 27.496, P < 0.001) which decreased when males developed at 35°C compared to those developing at 26°C (Figure 3). Overall, fertility decreased in both sexes when reared at 35°C, with the largest impacts on fecundity in females and fertility proportions and egg hatch proportions in males.

### Elevated temperatures during both development and adulthood reduce fertility

To determine the effect of elevated temperatures during adulthood on the fertility of both sexes, we performed a temperature shift experiment where mosquitoes developing at 26°C and 35°C were exposed to each temperature during adulthood. When we considered reproductive success, we found significant effects of both developmental temperature and adult temperature on the proportion of females laying eggs, the proportion of females with some viable eggs, and the proportion of females with some viable offspring (Two-way ANOVA: all P < 0.001). These effects were mainly driven by the low proportion of females laying eggs or producing viable offspring when male and female mosquitoes were kept at 35°C during development and adulthood (Figure 5).

**Figure 5.**
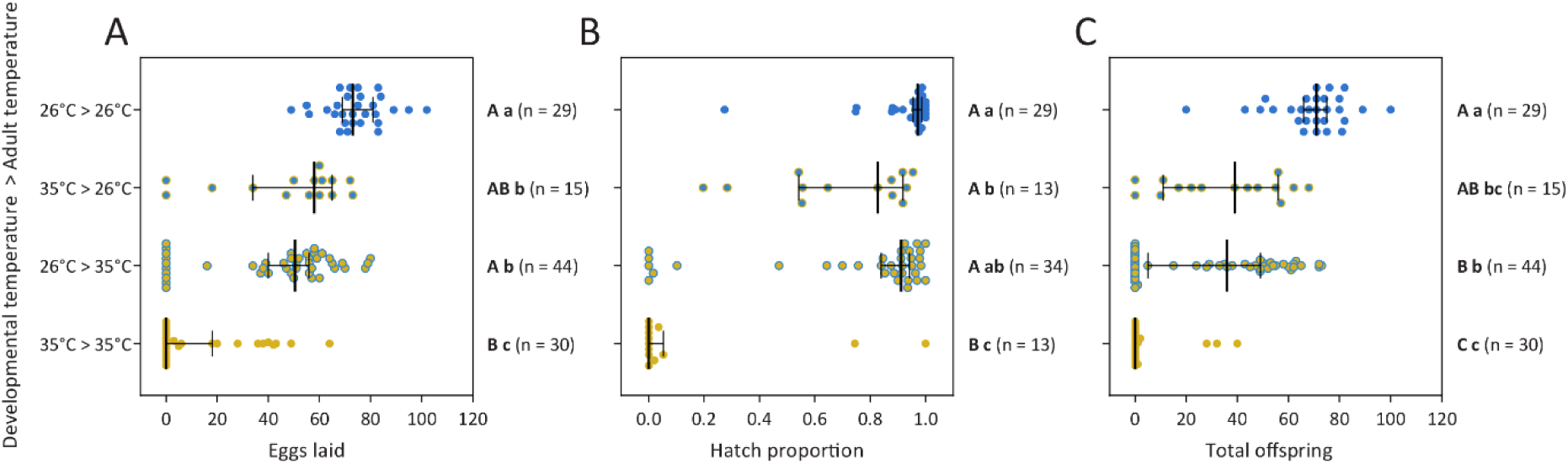
Effects of elevated developmental and adult temperatures on *Aedes aegypti* fertility. We measured (A) fecundity, (B) egg hatch proportions and (C) total offspring of mosquitoes reared at constant temperatures of 26 and 35°C until adult emergence then maintained at 26 or 35°C until laying eggs. Both sexes within a treatment were reared and maintained under the same conditions. Dots and error bars represent medians and 95% confidence intervals. Different letters next to plots indicate significant differences (P < 0.05) between treatments according to Tukey’s post-hoc tests, with capital letters for the binomial analyses of reproductive success and lowercase letters for the analyses of counts excluding zero values. The n values represent the number of replicates measured per treatment and trait.

In an analysis of the counts with zero values excluded, we found substantial effects of developmental temperature (two-way ANOVA, fecundity: F_1,85_ = 16.526, P < 0.001, hatch proportion: F_1,73_ = 24.316, P < 0.001, total offspring: F_1,73_ = 24.316, P < 0.001) on fertility (Figure 5), where rearing mosquitoes at 35°C decreased fertility consistent with the previous experiments. We also observed impacts of adult temperature, with maintenance at 35°C during adulthood significantly decreasing fecundity (F_1,85_ = 31.420, P < 0.001) and total offspring (F_1,73_ = 23.789, P < 0.001) but not egg hatch proportions (F_1,73_ = 2.539, P = 0.115). There were no significant interactions between developmental and adult temperature for any trait (all P > 0.105). The effects of elevated temperatures appeared to be cumulative across life stages, with mosquitoes maintained at 35°C during both development and adulthood producing a median of zero offspring (Figure 5).

### Elevated developmental temperatures improve fecundity in the subsequent generation

To examine the effects of elevated developmental temperatures across generations, we reared parental mosquitoes at 26°C or 35°C and then reared their offspring at each of these temperatures. As expected, offspring reared at 35°C experienced decreased fertility (Figure 6), with a signficiant effect of offspring rearing temperature on fecundity (two-way ANOVA: F_1,198_ = 126.492, P < 0.001), egg hatch (F_1,183_ = 52.794, P < 0.001) and total offspring (F_1,183_ = 155.528, P < 0.001) on the count data with zeros excluded. For the proportional data, offspring reared at 35°C had a significantly higher proportion with no eggs hatching (χ^2^ = 7.942, df = 1, P = 0.005) and a higher proportion with no viable offspring (χ^2^ = 12.069, df = 1, P < 0.001), but there was a no significant effect of offspring developmental temperature on the proportion of mosquitoes laying eggs (χ^2^ = 2.962, df = 1, P = 0.085). We also identified effects of parental rearing temperature on fecundity (F_1,198_ = 21.730, P < 0.001) and total offspring (F_1,183_ = 15.240, P < 0.001) but not egg hatch (F_1,192_ = 0.0001, P = 0.992). Notably, offspring from parents reared at 35°C showed increased fecundity at both rearing temperatures. However, there was no significant effect of parental developmental temperature on the proportion of females laying eggs, the proportion with viable eggs, or the proportion with viable offspring (all P > 0.535).

**Figure 6.**
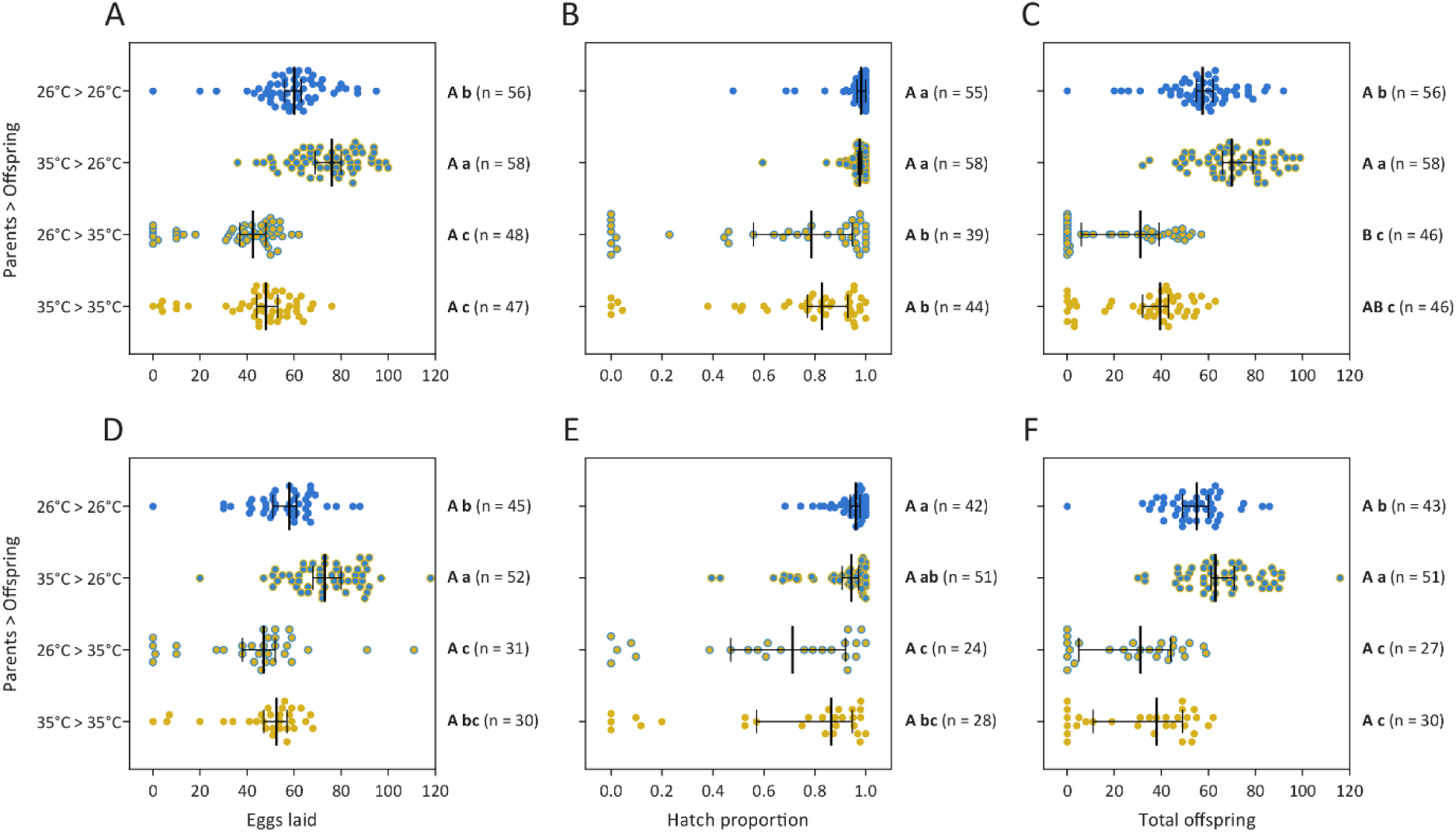
Cross-generational effects of elevated developmental temperatures on *Aedes aegypti* fertility persist across two gonotrophic cycles. We measured fertility in offspring reared at 26 or 35°C following rearing at 26 or 35°C in the parental generation. Both sexes within a treatment were reared at the same temperature. We measured (A, D) fecundity, (B, E) egg hatch proportions and (C, F) total offspring across the (A-C) first and (D-F) second gonotrophic cycles. Dots and error bars represent medians and 95% confidence intervals. Different letters next to plots indicate significant differences (P < 0.05) between treatments according to Tukey’s post-hoc tests, with capital letters for the binomial analyses of reproductive success and lowercase letters for the analyses of counts excluding zero values. The n values represent the number of replicates measured per treatment and trait.

### Effects of elevated developmental temperature on fertility persist across gonotrophic cycles

To assess whether the costs and benefits of elevated developmental temperatures persist into the second gonotrophic cycle, females from the previous experiment were blood-fed again. Consistent with the first gonotrophic cycle, developing at 35°C imposed significant costs in the second generation, with reduced fecundity (two-way ANOVA: F_1,149_ = 41.010, P < 0.001), egg hatch (F_1,136_ = 28.401, P < 0.001) and total offspring (F_1,136_ = 72.298, P < 0.001, Figure 6). Costs also persisted for reproductive success in terms of the proportion of females with no eggs hatching (χ^2^ = 6.917, df = 1, P = 0.009) and the proportion with no viable offspring (χ^2^ = 10.704, df = 1, P = 0.001). Across all individuals, there were signficant correlations between the first and second gonotrophic cycles for fecundity (Spearman r = 0.564, P < 0.001), egg hatch (r = 0.497, P < 0.001) and total offspring (r = 0.680, P < 0.001, Figure S1). Parental effects on offspring fertility were also apparent in the second gonotrophic cycle, where elevated temperatures experienced by the parents led to increased fecundity (F_1,149_ = 16.446, P < 0.001) and total offspring (F_1,136_ = 12.159, P < 0.001) but not egg hatch proportions (F_1,136_ < 0.001, P = 0.997, Figure 6). Consistent with the first gonotrophic cycle, there was no significant effect of parental developmental temperature on reproductive success (all P > 0.444).

## Discussion

Our study has identified opposing effects of elevated developmental temperatures on *Ae. aegypti* fitness. While mosquito reproductive success and fertility is reduced below the developmental lethal thermal limit, particularly when combined with elevated temperatures during adulthood, this is offset by an increase in CTmax and beneficial cross-generational effects on fecundity. The costs to fertility occur below the egg to adult viability limit (LT_50_ of 35.51°C), with the traits affected being sex-specific. Consequently, these factors may limit the persistence of *Ae. aegypti* in certain climatic conditions. Importantly, our work demonstrates how measuring fecundity alone may underestimate the effects of heat stress on male fertility as females may lay non-viable eggs.

It is well established that the fertility of females and males can differ in their heat sensitivity (Iossa, 2019; Ørsted et al., 2024; Parrett et al., 2024; Weaving et al., 2024). While past studies have demonstrated costs of elevated developmental temperatures to mosquito fertility, our study is the first to delineate these effects by sex. Warmer temperatures below the lethal limit resulted in decreased fecundity in females. This finding aligns with earlier research indicating that female body size (Farjana et al., 2012; Mohammed & Chadee, 2011) and ovariole numbers (Bader & Williams, 2012) decrease with rising temperatures. In contrast to females, the reduction in overall fertility observed in males subjected to elevated developmental temperatures was largely driven by costs to hatch proportions. In certain cases, there was also an increase in the number of females laying no eggs, potentially reflecting diminished mating success or reduced sperm quantity and quality due to decreased body size (Felipe Ramirez-Sanchez et al., 2020; Ponlawat & Harrington, 2007, 2009) or the direct impacts of elevated temperatures on spermatogenesis (Canal Domenech & Fricke, 2023; Gandara & Drummond-Barbosa, 2023). Notably, the impacts on both fecundity and egg viability persisted across two gonotrophic cycles suggesting that females are unable to recover from costs expressed in both males and females, while also indicating an absence of delayed costs. While our crosses were only performed at a single point post-emergence, the timing of mating is an important consideration for future research, given that both male (Canal Domenech & Fricke, 2022) and female (Walsh et al., 2022) fertility can recover through remating in *Drosophila*.

Our study highlights additive costs of elevated temperatures during development and adulthood to fertility. At a constant temperature of 35°C, fertilility was reduced to a median of zero, falling below the egg to adult viability LT_50_ of 35.5°C. This finding adds to recent literature documenting ferility thermal limits lower than developmental and adult lethal limits in *Drosophila* and other species (Parratt et al., 2021; Parrett et al., 2024; Van Heerwaarden & Sgrò, 2021). While we did not assess adult longevity in this study, it is likely that elevated temperatures at both immature and adult stages further decrease fitness beyond effects on fertility, as demonstrated in other mosquito species (Christiansen-Jucht et al., 2014; Ezeakacha & Yee, 2019). Elevated temperatures during adulthood also affect other aspects of mosquito biology, including host-seeking (Lahondère et al., 2023) and blood feeding success (Christiansen-Jucht et al., 2015) which were not considered in our experiments. Our methodology, which involved transferring mosquitoes between temperatures at emergence and prior to mating, limited our ability to determine whether the costs of elevated temperatures during adulthood were sex-specific or driven by temperatures before, during or after mating. Previous studies have documented the effects of elevated adult temperatures on female *Ae. aegypti,* revealing that oviposition can be delayed or completely inhibited at high temperatures during adulthood (Carrington, Armijos, et al., 2013; Costa et al., 2010). Although copulation and insemination appear unaffected at 35°C in *Ae. aegypti* (Bader & Williams, 2012), elevated pre-mating temperatures have been shown to reduce insemination frequencies in other mosquito species (Horsfall & Taylor, 1967), leaving the resulting impacts on female fertility unclear.

The positive effects of elevated developmental temperatures we identified were relatively limited compared to the costs described above. For CTmax, ARRs around 0.05°C were similar to those reported for *Drosophila melanogaster* (ARR 0.030-0.068 van Heerwaarden et al. (2016), but substantially lower than in other mosquito species within a similar temperature range (0.22 in *Ae. aegypti* males (Sasmita et al., 2019), 0.11 in *Culex pipiens* females (Gray, 2013), 0.14 in *Anopheles funestus* (Lyons et al., 2012) and 0.20 in *Anopheles arabiensis* (Lyons et al., 2012)). The cross-generational benefits to fertility were more substantial, with a 12-32% increase in total offspring for females reared at 35°C in the parental generation (depending on the gonotrophic cycle and offspring rearing temperature). However, since developmental temperatures of 35°C did induce mortality, we are unable to separate whether effects reflect plastic responses or genetic changes in the population due to selection. Nonetheless, the results here suggest that beneficial acclimation and the accumulation of heat damage across life stages, both within and across generations, are likely to impact survival and fertility in complex ways that may not be easily predicted from damage models alone (Jørgensen et al., 2021; Ørsted et al., 2022; Rezende et al., 2020).

We acknowedge several limitations in our study design that could influence our estimates of elevated temperature effects. Although our experiments utilized a single laboratory population, mosquitoes show intraspecific variation in heat tolerance (Chakraborty et al., 2024; Dennington et al., 2024; Vorhees et al., 2013), cold tolerance (De Majo et al., 2019; Kramer et al., 2021; Zani et al., 2005), adult desiccation tolerance (Ross et al., 2023) and quiescent egg viability (Faull & Williams, 2015). Responses in our study could also be influenced by laboratory adaptation given that heat tolerance may differ between near-field and established mosquito populations (Ciota et al., 2014; Dennington et al., 2024). Furthermore, our crossing design, which relied on group matings rather than individual pairings, could influence fertility estimates. While females typically mate once (Degner & Harrington, 2016), males can inseminate multiple females in a short period, where sperm depletion in multiply mated males may affect female fertility (Ponlawat and Harrington 2007, Ramirez-Sanchez et al. 2020). While single pair crosses are often used to investigate temperature effects on reproduction (Evans et al., 2018; Snook et al., 2000), logistical challenges make such designs difficult for *Ae. aegypti*, as they require more space for mating. Moreover, single-pair crosses may not accurately represent field conditions, where males typically mate with multiple females.

Our results have important implications for models predicting future mosquito distributions under climate change. Many models treat species as uniform and unchanging entities (Campbell et al., 2015; Kraemer et al., 2019; Messina et al., 2019), but incorporating both plastic responses and evolutionary changes can significantly influence predictions of species distribution and abundance (Brass et al., 2024; Bush et al., 2016; Kearney et al., 2009; Valladares et al., 2014). We suggest that sub-lethal impacts on fertility, which occur at temperatures below the lethal limit, should also be considered. However, estimating these impacts under natural conditions remains challenging.

Additionally, while we only investigated constant temperatures in this study, temperature fluctuations can have significant effects on fitness, even when mean temperatures are identical (Carrington, Seifert, Willits, et al., 2013). Larval habitats are highly variable in their thermal profiles across space and time, and mosquitoes are likely exposed to different combinations of thermal stress at various life stages. High temperatures during the egg and adult stages, in combination with low humidity, can further increase mortality due to desiccation (Brown et al., 2023). These effects may be further exacerbated by resource competition, which can extend development (Couret et al., 2014), potentially interacting with temperature to reduce fitness through cumulative heat damage (Huxley et al., 2021; Padmanabha et al., 2011).

## Acknowledgements

We thank Apeksha Warusawithana, Ella Yeatman and Mel Berran for technical assistance with experiments. We also thank Fernando Gabriel Noriega for providing helpful comments on the manuscript.

## Supporting information

**Figure S1.**
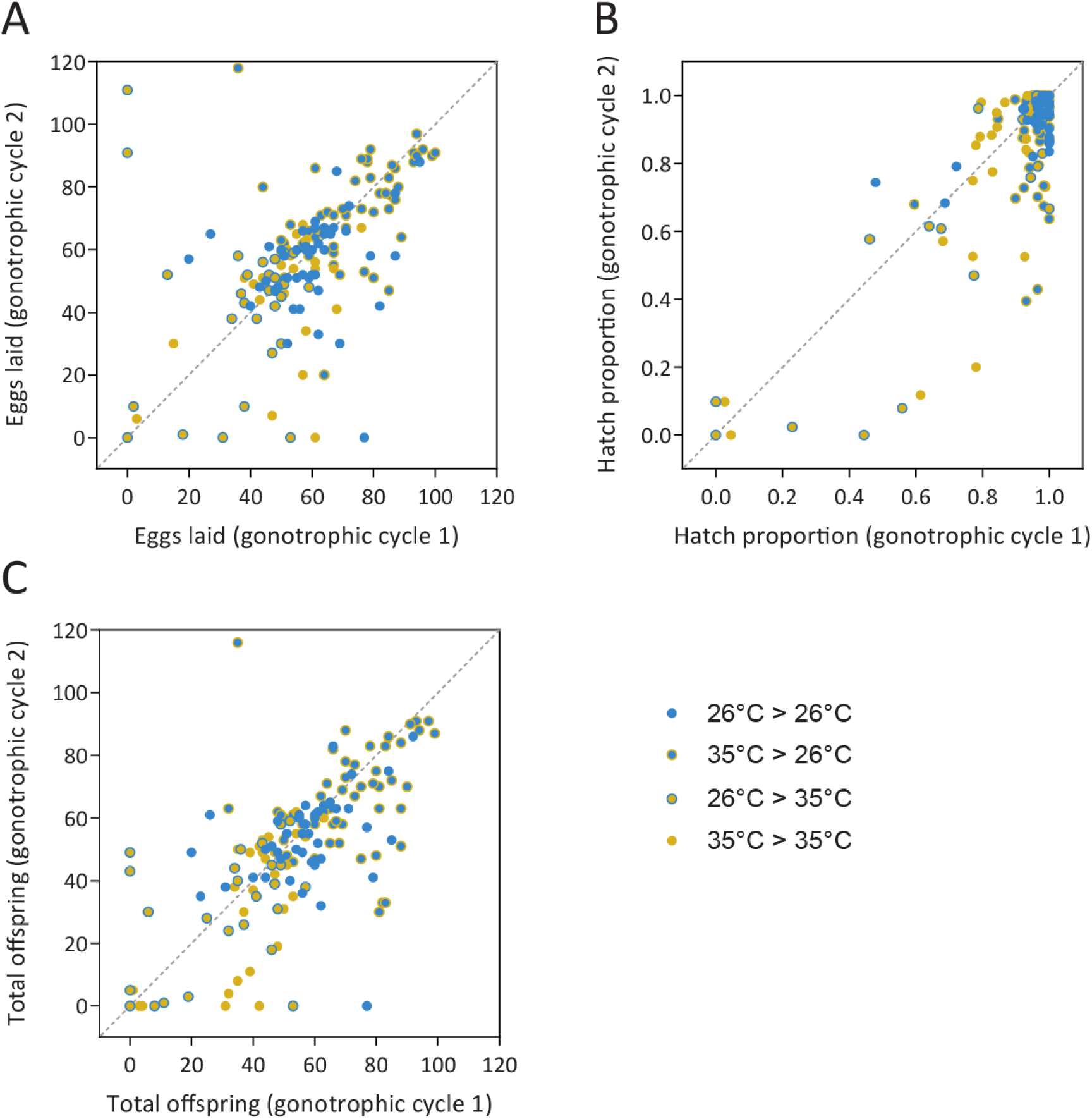
Correlations in (A) fecundity, (B) egg hatch proportions and (C) total offspring between the first and second gonotrophic cycles. Dots represent data from individual females measured across two gonotrophic cycles in the experiment testing cross-generational effects of elevated developmental temperatures also shown in Figure 6.

